# Analysis of changes occurring in Codon Positions due to mutations through the cellular automata transition rules

**DOI:** 10.1101/2021.08.30.458305

**Authors:** Antara Sengupta, Sreeya Ghosh, Pabitra Pal Choudhury

## Abstract

Variation in the nucleotides of a codon may cause variations in the evolutionary patterns of a DNA or amino acid sequence. To address the capability of each position of a codon to have non-synonymous mutations, the concept of degree of mutation has been introduced. The degree of mutation of a particular position of codon defines the number of non-synonymous mutations occurring for the substitution of nucleotides at each position of a codon, when other two positions of that codon remain unaltered. A Cellular Automaton (CA), is used as a tool to model the mutations of any one of the four DNA bases A, C, T and G at a time where the DNA bases correspond to the states of the CA cells. Point mutation (substitution type) of a codon which characterizes changes in the amino acids, have been associated with local transition rules of a CA. Though there can be 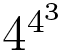 transitions of a 4-state CA with 3-neighbourhood cells, here it has been possible to represent all possible point mutations of a codon in terms of combinations of 16 local transition functions of the CA. Further these rules are divided into 4 classes of equivalence. Also, according to the nature of mutations, the 16 local CA rules of substitutions are classified into 3 sets namely, ‘No Mutation’, ‘Transition’ and ‘Transversion’. The experiment has been carried out with three sets of single nucleotide variations(SNVs) of three different viruses but the symptoms of the diseases caused by them are to some extent similar to each other. They are SARS-CoV-1, SARS-CoV-2 and H1N1 Type A viruses. The aim is to understand the impact of nucleotide substitutions at different positions of a codon with respect to a particular disease phenotype.

## 1. Introduction

Genetic code defines some rules to translate genetic information encoded in nucleotide triplets or codons into amino acids. It also defines the order of amino acid to be added next during protein synthesis. 4^3^ = 64 codons are there in genetic code table, which encodes 20 standard amino acids and 3 stop codons. Hence, there arises a context of degeneracy. Multiplet structure of DNA sequence [1] specifies that instead of one-to-one mapping a single amino acid can be coded by one, two, three, four or six codons. The codon usage is an important determinant of gene expression and surprisingly transcriptions rather than translations play a key role here [2, 3]. It has been reported that instead of codons or amino acids, codon and amino acid usage is consistent with the forces acting on four DNA bases [4]. Analysis of codon usage gives insight about the evolution of any organism [5]. Selection of codon to code for an amino acid is a natural selection and amino acid composition in protein aims to minimize the the impact of mutations on protein structure [6]. A codon can have mutations at the first, second or third positions. Mutations at the third position of the codon are more likely to be synonymous than mutations that occur at the first or second positions [7]. Hence, probability of substitution of amino acid with a new one due to mutations at third position of a codon is less than that of its first and second positions. The second position of codon is the most conserved position, as every nucleotide change in this position leads to substitution of another amino acid [8]. Hence, change of nucleotides at a particular position of a codon due to substitutions have positional impact on the change in amino acids.

Researchers throughout the globe are trying to figure out the pattern of mutations responsible for a particular genetic disease. Numerous mathematical model based approaches are already introduced to make quantitative understanding of a disease and to apply classification rules to segregate that disease from others. Now-a-days when some Asian countries are witnessing 2nd wave of Corona and 3rd wave has already arrived in other continents like Europe, researchers are trying in every way to understand the pattern of mutations taking place in SARS-CoV-2. A Plethora of mathematical models are already introduced to reach to that goal [9, 10]. Scientists are working over differentiating coronavirus from influenza virus as both the disease COVID-19 and flu have some similar type of symptoms [11]. Mutations may lead to occur biodiversity. Biodiversity is characterized by the continual replacement of branches in the tree of life, that is clade [12]. People are trying to reach the origin of the tree of life to get some ways of prevention [13].

Cellular Automaton (pl. cellular automata, abbrev. CA) [14] is a discrete model introduced by J.von Neumann and S.Ulam in 1940s for designing self replicating systems. It consists of a finite/countably infinite number of finite-state semi-automata known as ‘cells’ arranged in an ordered *n*-dimensional grid. Each cell receives input from the neighbouring cells and changes according to a transition function. Application of cellular automata in bioinformatics is a well-known approach [15]. There has been studies on the evolution of DNA sequence using automata [16], and CA transition rules [17, 18]. CA based models are used to unfold different facts in genomics, proteomics [19, 20] and even for the representation of protein translation using CA rules [21]. However representing all possible point mutations of a codon in terms of CA rules have not been addressed earlier. Variation in mutations and codon selection may cause differences in evolutionary patterns across a DNA or amino acid sequence [7]. In this present study, we have been able to represent all possible changes in amino acids due to point mutations of codons in terms of combinations of 16 local transition rules of a CA. These could further be divided into 4 classes of equivalence. Depending upon the capability of producing a new amino acid, degree of mutations of codons at 3 different positions have been derived. Also, according to the nature of mutations, the 16 local CA rules of substitutions are classified here into 3 sets namely, ‘No Mutation’, ‘Transition’ and ‘Transversion’.

Recently, attempts has been made to model COVID-19 spread within the framework of Probabilistic CA [22] and Fuzzy CA [23]. Pokkuluri et.al. [24] have constructed CA based classifiers to predict the trend of SARS-CoV-2. Few papers are reported, where dynamics of the influenza infection is described using Beauchemin’s CA model [25, 26, 27]. In our work, we have considered SNVs of three different viruses manifesting similar symptoms, namely SARS-CoV-1, SARS-CoV-2 and H1N1 Type A viruses. Our objective is to get a pattern of mutations occurring in these diseases, in the light of degrees of mutations and CA transition functions.

## 2. Methods and Materials

### 2.1. Derive Degree of Mutation of Nucleotides at Different Positions of Codon

According to the genetic code table 61 codons code for 20 amino acids and there are three stop codons [28]. The standard classic model of genetic code table consists four rows and four columns. The four rows represents the first base of each codon, the four columns represent the second base and the right side indicates the third base of them. Codon contains combinations of 4 bases A,T,C,G at its 3 positions and as a whole codes for a particular amino acid. Since there are 20 different amino acids and 64 possible codons, more than one codon may code for a single amino acid. Hence, any changes in nucleotides at any positions of codon due to mutation either may change the produced amino acid or can code for the same amino acid and there is a talk about non-synonymous and synonymous mutations respectively. Here in this section it is tried to get a clear view of mapping between codon and amino acid when mutations occur at first, second and third positions of a codon.

#### Definition 2.1

(Degree of Mutation of a particular position of codon). *Given a codon C with constituent nucleotides say*, (*N*_1_, *N*_2_, *N*_3_), *where N*_i_ ∈ *N is a particular position of a codon. Now, consider S_i_ as any one nucleotide among the set of nucleotides S={T,C,A,G} at a particular position N_i_ in codon C when nucleotides at other two positions are constant. The degree of mutation (δ(M)) at a particular position of codon defines the number of non-synonymous mutations occurred to substitution of nucleotides at that position of a codon, when other two positions of that codon are unaltered*.

It is to be noted that when any two positions of a codon are constant, it is possible to make change in 16 possible places with maximum three nucleotides, when the position is initially being occupied by any one of four nucleotides (shown in Table 1). Thus number of probable changes of amino acids (AA) due to mutations at first position of codon may be between 0 to 3 when 2nd and 3rd positions are constant or fixed. Hence, the degree of mutation at the first position of codon vary from 0 to 3. Due to mutation change in nucleotide at 2nd position the probable change in amino acid will be maximum and the degree of mutation is between the range of 2 to 3. The mutations at third position of codon have very less capability to make non-synonymous changes in amino acids and hence the range is 0 to 2. As an example, when T is constant at both 2nd and 3rd positions, due to change in nucleotides (A/T/C/G) at first position the total numbers of amino acids can be changed is 3 according to genetic code table and the amino acids are F, L, I and V. Hence, the degree of mutation *δ*(*M*) here is 3.

**Table 1:** Degree of mutation of all 64 codons

| 1st Position | 2nd Position | 3rd Position | $\delta(M)$ at 1st position | Name of AA |
| --- | --- | --- | --- | --- |
| T/C/A/G | T | T | 3 | F/L/I/V |
| T/C/A/G | T | C | 3 | F/L/I/V |
| T/C/A/G | T | A | 2 | L/M/V |
| T/C/A/G | T | G | 2 | L/M/V |
| T/C/A/G | C | T | 3 | S/P/T/A |
| T/C/A/G | C | C | 3 | S/P/T/A |
| T/C/A/G | C | A | 3 | S/P/T/A |
| T/C/A/G | C | G | 3 | S/P/T/A |
| T/C/A/G | A | T | 3 | Y/H/N/D |
| T/C/A/G | A | C | 3 | Y/H/N/D |
| T/C/A/G | A | A | 3 | STOP CODON/Q/K/E |
| T/C/A/G | A | G | 3 | STOP CODON/Q/K/E |
| T/C/A/G | G | T | 3 | C/R/S/G |
| T/C/A/G | G | C | 3 | C/R/S/G |
| T/C/A/G | G | A | 2 | STOP CODON/R/R/G |
| T/C/A/G | G | G | 2 | W/R/R/G |
| 1st Position | 2nd Position | 3rd Position | $\delta(M)$ at 2nd position | Name of AA |
| T | T/C/A/G | T | 3 | F/S/Y/C |
| T | T/C/A/G | C | 3 | F/S/Y/C |
| T | T/C/A/G | A | 2 | L/S/STOP CODON/STOP CODON |
| T | T/C/A/G | G | 3 | L/S/STOP CODON/W |
| C | T/C/A/G | T | 3 | L/P/H/R |
| C | T/C/A/G | C | 3 | L/P/H/R |
| C | T/C/A/G | A | 3 | L/P/Q/R |
| C | T/C/A/G | G | 3 | L/P/Q/R |
| A | T/C/A/G | T | 3 | I/T/N/S |
| A | T/C/A/G | C | 3 | I/T/N/S |
| A | T/C/A/G | A | 3 | I/T/K/R |
| A | T/C/A/G | G | 3 | I/T/K/R |
| G | T/C/A/G | T | 3 | V/A/D/G |
| G | T/C/A/G | C | 3 | V/A/D/G |
| G | T/C/A/G | A | 3 | V/A/E/G |
| G | T/C/A/G | G | 3 | V/A/E/G |
| 1st Position | 2nd Position | 3rd Position | $\delta(M)$ at 3rd position | Name of AA |
| T | T | T/C/A/G | 1 | F/L |
| T | C | T/C/A/G | 0 | S |
| T | A | T/C/A/G | 1 | Y/STOP CODON |
| T | G | T/C/A/G | 2 | C/STOP CODON/W |
| C | T | T/C/A/G | 0 | L |
| C | C | T/C/A/G | 0 | P |
| C | A | T/C/A/G | 1 | H/Q |
| C | G | T/C/A/G | 0 | R |
| A | T | T/C/A/G | 1 | I/M |
| A | C | T/C/A/G | 0 | T |
| A | A | T/C/A/G | 1 | N/K |
| A | G | T/C/A/G | 1 | S/R |
| G | T | T/C/A/G | 0 | V |
| G | C | T/C/A/G | 0 | A |
| G | A | T/C/A/G | 1 | D/E |
| G | G | T/C/A/G | 0 | G |

### 2.2. Cellular Automata and Mutations of Nucleotides

#### Definition 2.1.

*A **CA**(denoted by* 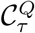*)(reported in [29, 14, 30]) is a triplet (Q*,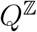,*τ), where*,

- *Q is a finite state set*
- 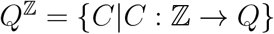 *is the set of all global configurations C*
- 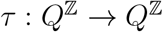 *is a global transition function*

#### Definition 2.2.

*A restriction from* 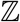 *to a subset S_i_ containing* 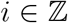, *induces a restriction of C to* 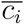 *given by* 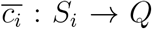; *where* 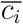 *may be called the **local configuration** and S_i_ the **neighbourhood** of the i^th^ cell*.

*The mapping μ_i_: Q^S_i_^ → Q is known as a **local transition function** for the i^th^ cell*.

*Thus* 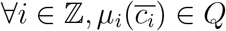 *and it follows that*,

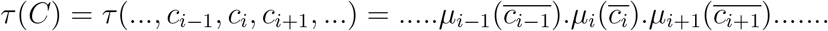

### 2.3. Representation of Mutations Occurring in Different Codon Positions Using CA Transitions

Point mutation of a codon can be associated with local transitions of Cellular Automata (CA) having 3-celled local configurations. 16 substitutions are possible with four DNA bases A, C, T and G. They are

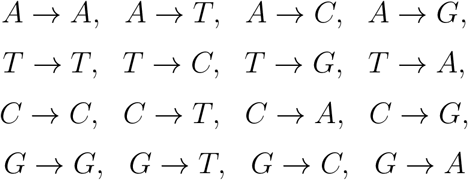

They can be represented in terms of combinations of 16 local transition functions of CA.

Let us consider the global configuration of a CA to be composed of local configurations having three cells corresponding to the three nucleotide positions of a codon. The position of the codon at which the point mutation occurs, is denoted by the *i^th^* cell and the other two nucleotides which remain fixed are denoted by *x* and *y* where *x*, *y* ∈ *Q* = {*A*, *T*, *C*, *G*}. If point mutation occurs at the third position then the neighbourhood of the *i^th^* cell is considered as

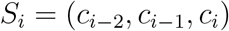

The local configuration of the *i^th^* cell maybe denoted by (*x*, *y*, *c_i_*) such that *c*_*i*-2_ = *x* and *c*_*i*-2_ = *y*. The local transition function for *i^th^* cell denoted by *μ*_*R*_(*xyi*)__ is,

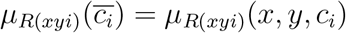

where *R*(*xyi*) is the rule number for some *R* ∈ {0,1,2,…, 15}. The rules for third position mutation corresponding to first and second position constant nuleotides *x*, *y* is computed as follows:

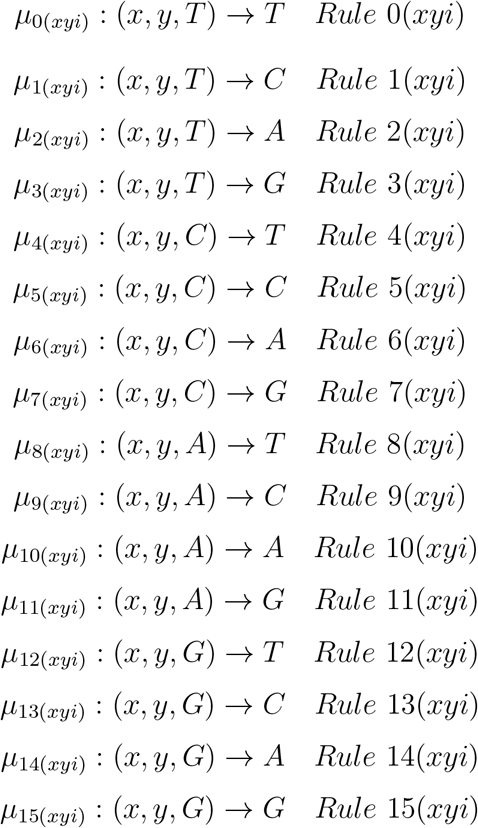

#### Example 2.1.

*For constant first and second necleotides AA, Rule* 6(*AAi*) *represented by μ*_6_(*AAi*)__: (*A*, *A*, *C*) → *A changes nucleotide C in the third position to nucleotide A corresponding to the mutation of codon AAC to AAA for amino acid Asn to Lys*.

Substitutions at second and first postions can be computed similar to that of the third position point mutation by changing the neighbourhood of the *i^th^* cell as follows.

#### Corollary 2.1.

*If point mutation occurs at the second position then the neighbourhood of the i^th^ cell is considered as*

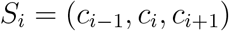

*The local configuration of the i^th^ cell maybe denoted by* (*x*, *c_i_*, *y*) *such that c*_*i*-1_ = *x and c*_*i*+1_ = *y. The local transition function for i^th^ cell denoted by μ_R_(xiy)__ is*,

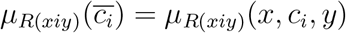

*The rules R(xiy) for second position mutation are as follows*:

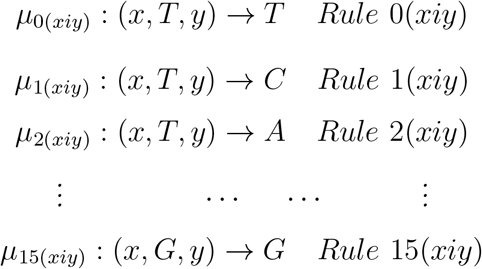

#### Corollary 2.2.

*If point mutation occurs at the first position then the neighbourhood of the i^th^ cell is considered as*

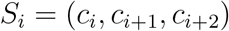

*The local configuration of the i^th^ cell maybe denoted by (c_i_, x, y) such that c*_*i*+1_ = *x and c*_*i*+2_ = *y. The local transition function for i^th^ cell denoted by μ_R_(xyi)__ is*,

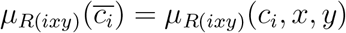

*The rules for first position mutation are as follows*:

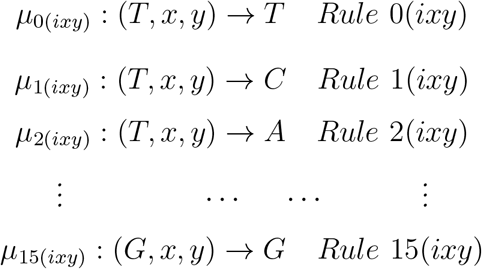

These combinations of 16 CA rules can further be classified into three sets which depict No Mutation, Transition and Transversion of nucleotides irrespective of the position where the point mutation occurs. According to the rules for point mutation with respect to constant nucleotides x and y we get:

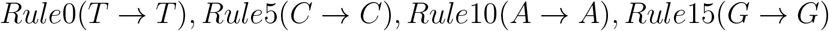

representing No Mutations;

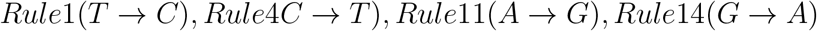

representing Transitions where point mutations occur due to substitutions between any two purine (A or G) bases or pyrimidine bases (T or C);

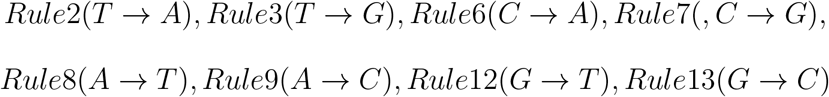

representing Transversions where point mutations occur due to substitution of a purine (A or G) base by a pyrimidine base (T or C) or vice-versa.

These classifications have been tabulated in Table 2.

**Table 2:** Classification of CA rules

| CA RULES | CLASSIFICATION |
| --- | --- |
| 0($T \rightarrow T$)<br>5($C \rightarrow C$)<br>10($A \rightarrow A$)<br>15($G \rightarrow G$) | NO MUTATION |
| 1($T \rightarrow C$)<br>4($C \rightarrow T$)<br>11($A \rightarrow G$)<br>14($G \rightarrow A$) | TRANSITION |
| 2($T \rightarrow A$)<br>3($T \rightarrow G$)<br>6($C \rightarrow A$)<br>7($C \rightarrow G$)<br>8($A \rightarrow T$)<br>9($A \rightarrow C$)<br>12($G \rightarrow T$)<br>13($G \rightarrow C$) | TRANSVERSION |

### 2.4. Amino Acids Arising due to Point Mutations Represented by Equivalent Rules

#### Definition 2.3.

*Any two local transition functions for an i^th^ cell denoted by μ_R_(xyi)__ and μ_R′_(xyi)__ are equivalent if both the rules produce same output. Thus*

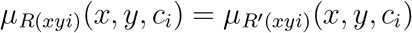

*where R(xyi) and R′(xyi) are rule numbers for R, R′* ∈ {0, 1, 2,…, 15}.

Any two equivalent rules belong to same class of 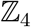 where 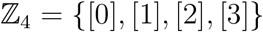.

From the list of rules we get that, if point mutation occurs at the third position then:

1. [0](*xyi*) = {*Rule* 0(*xyi*), *Rule* 4(*xyi*), *Rule* 8(*xyi*), *Rule* 12(*xyi*)}
2. [1](*xyi*) = {*Rule* 1(*xyi*), *Rule* 5(*xyi*), *Rule* 9(*xyi*), *Rule* 13(*xyi*)}
3. [2](*xyi*) = {*Rule* 2(*xyi*), *Rule* 6(*xyi*), *Rule* 10(*xyi*), *Rule* 14(*xyi*)}
4. [3](*xyi*) = {*Rule* 3(*xyi*), *Rule* 7(*xyi*), *Rule* 11(*xyi*), *Rule* 15(*xyi*)}

Correspondingly, amino acids produced from codon having nucleotide base

1. T in the 3rd position is obtained by applying [0](*xyi*)
2. C in the 3rd position is obtained by applying [0](*xyi*)
3. A in the 3rd position is obtained by applying [0](*xyi*)
4. G in the 3rd position is obtained by applying [0](*xyi*)

If point mutation (substitutions) occurs at second or first position then similar rules are equivalent. Also amino acids produced from corresponding point mutations of codons can be obtained similarly. Thus the table 3 shows all possible changes in codons due to point mutations and possible changes in amino acids due to it through the light of CA rules.

**Table 3:** CA rules to identify changes in codons due to point mutations and possible changes in amino acids due to it

| Amino acid | Codon | Point mutation | Fixed nucleotides | Rules |
| --- | --- | --- | --- | --- |
| Phe | TTT/TTC | 3rd<br>2nd<br>1st | TT<br>TT/TC<br>TT/TC | [0](TTi), [1](TTi)<br>[0](TiT), [0](TiC)<br>[0](iTt), [1](iTc) |
| Leu | TTA/TTG | 3rd<br>2nd<br>1st | TT<br>TA/TG<br>TA/TG | [2](TTi), [3](TTi)<br>[0](TiA), [0](TiG)<br>[0](iTa), [0](iTg) |
|  | CTT/CTC/CTA/CTG | 3rd<br>2nd<br>1st | CT<br>CT/CC/CA/CG<br>TT/TC/TA/TG | [0](CTi), [1](CTi), [2](CTi), [3](CTi)<br>[0](CiT), [0](CiC), [0](CiA), [0](CiG)<br>[1](iTt), [1](iTc), [1](iTa), [1](iTg) |
| Ile | ATT/ATC/ATA | 3rd<br>2nd<br>1st | AT<br>AT/AC/AA<br>TT/TC/TA | [0](ATi), [1](ATi), [2](ATi)<br>[0](AiT), [0](AiC), [0](AiA)<br>[2](iTt), [2](iTc), [2](iTa) |
| Met | ATG | 3rd<br>2nd<br>1st | AT<br>AG<br>TG | [3](ATi)<br>[0](AiG)<br>[2](iTg) |
| Val | GTT/GTC/GTA/GTG | 3rd<br>2nd<br>1st | GT<br>GT/GC/GA/GG<br>TT/TC/TA/TG | [0](GTi), [1](GTi), [2](GTi), [3](GTi)<br>[0](GiT), [0](GiC), [0](GiA), [0](GiG)<br>[3](iTt), [3](iTc), [3](iTa), [3](iTg) |
| Ser | TCT/TCC/TCA/TCG | 3rd<br>2nd<br>1st | TC<br>TT/TC/TA/TG<br>CT/CC/CA/CG | [0](TCi), [1](TCi), [2](TCi), [3](TCi)<br>[1](TiT), [1](TiC), [1](TiA), [1](TiG)<br>[0](iCt), [0](iCc), [0](iCa), [0](iCg) |
|  | AGT/AGC | 3rd<br>2nd<br>1st | AG<br>AT/AC<br>GT/GC | [0](AGi), [1](AGi)<br>[3](AiT), [3](AiC)<br>[2](iGt), [2](iGc) |
| Pro | CCT/CCC/CCA/CCG | 3rd<br>2nd<br>1st | CC<br>CT/CC/CA/CG<br>CT/CC/CA/CG | [0](CCi), [1](CCi), [2](CCi), [3](CCi)<br>[1](CiT), [1](CiC), [1](CiA), [1](CiG)<br>[1](iCt), [1](iCc), [1](iCa), [1](iCg) |
| Thr | ACT/ACC/ACA/ACG | 3rd<br>2nd<br>1st | AC<br>AT/AC/AA/AG<br>CT/CC/CA/CG | [0](ACi), [1](ACi), [2](ACi), [3](ACi)<br>[1](AiT), [1](AiC), [1](AiA), [1](AiG)<br>[2](iCt), [2](iCc), [2](iCa), [2](iCg) |
| Ala | GCT/GCC/GCA/GCG | 3rd<br>2nd<br>1st | GC<br>GT/GC/GA/GG<br>CT/CC/CA/CG | [0](GCi), [1](GCi), [2](GCi), [3](GCi)<br>[1](GiT), [1](GiC), [2](GiA), [3](GiG)<br>[3](iCt), [3](iCc), [3](iCa), [3](iCg) |
| Tyr | TAT/TAC | 3rd<br>2nd<br>1st | TA<br>TT/TC<br>AT/AC | [0](TAi), [1](TAi)<br>[1](TiT), [1](TiC)<br>[0](iAt), [0](iAc) |
| His | CAT/CAC | 3rd<br>2nd<br>1st | CA<br>CT/CC<br>AT/AC | [0](CAi), [1](CAi)<br>[2](CiT), [2](CiC)<br>[1](iAt), [1](iAc) |
| Gln | CAA/CAG | 3rd<br>2nd<br>1st | CA<br>CA/CG<br>AA/AG | [2](CAi), [3](CAi)<br>[2](CiA), [2](CiG)<br>[1](iAA), [1](iAG) |
| Asn | AAT/AAC | 3rd<br>2nd<br>1st | AA<br>AT/AC<br>AT/AC | [0](AAi), [1](AAi)<br>[2](AiT), [2](AiC)<br>[2](iAt), [2](iAc) |
| Lys | AAA/AAG | 3rd<br>2nd<br>1st | AA<br>AA/AG<br>AA/AG | [2](AAi), [3](AAi)<br>[2](AiA), [2](AiG)<br>[2](iAA), [2](iAG) |
| Asp | GAT/GAC | 3rd<br>2nd<br>1st | GA<br>GT/GC<br>AT/AC | [0](GAi), [1](GAi)<br>[2](GiT), [2](GiC)<br>[3](iAt), [3](iAc) |
| Glu | GAA/GAG | 3rd<br>2nd<br>1st | GA<br>GA/GG<br>AA/AG | [2](GAi), [3](GAi)<br>[2](GiA), [2](GiG)<br>[3](iAA), [3](iAG) |
| Cys | TGT/TGC | 3rd<br>2nd<br>1st | TG<br>TT/TC<br>GT/GC | [0](TGi), [1](TGi)<br>[3](TiT), [3](TiC)<br>[0](iGt), [0](iGc) |
| Trp | TGG | 3rd<br>2nd<br>1st | TG<br>TG<br>GG | [3](TGi)<br>[3](TiG)<br>[0](iGg) |
| Arg | CGT/CGC/CGA/CGG | 3rd<br>2nd<br>1st | CG<br>CT/CC/CA/CG<br>GT/GC/GA/GG | [0](CGi), [1](CGi), [2](CGi), [3](CGi)<br>[3](CiT), [3](CiC), [3](CiA), [3](CiG)<br>[1](iGt), [1](iGc), [1](iGa), [1](iGg) |
|  | AGA/AGG | 3rd<br>2nd<br>1st | AG<br>AA/AG<br>GA/GG | [2](AGi), [3](AGi)<br>[3](AiA), [3](AiG)<br>[2](iGA), [2](iGG) |
| Gly | GGT/GGC/GGA/GGG | 3rd<br>2nd<br>1st | GG<br>GT/GC/GA/GG<br>GT/GC/GA/GG | [0](GGi), [1](GGi), [2](GGi), [3](GGi)<br>[3](GiT), [3](GiC), [3](GiA), [3](GiG)<br>[3](iGt), [3](iGc), [3](iGA), [3](iGG) |
| Stop Codon | TAA/TAG | 3rd<br>2nd<br>1st | TA<br>TA/TG<br>AA/AG | [2](TAi), [3](TAi)<br>[2](TiA), [2](TiG)<br>[0](iAA), [0](iAG) |
|  | TGA | 3rd<br>2nd<br>1st | TG<br>TA<br>GA | [2](TGi)<br>[3](TiA)<br>[0](iGA) |

**Table 4:** Dataset Specification

| Gene Name | # Isolates | # SNVs |
| --- | --- | --- |
| SARS-CoV-2 | 39680 | 85 |
| SARS-CoV-1 | 54 | 342 |
| H1N1 | 35008 | 8 |

## 3. Results and Discussion

### 3.0.1. Collection of Genomic Sequences

To establish the novelty of the methodologies discussed in previous section, it is necessary to apply the same into a given dataset. To carry out the experiment, mutated genomic sequences of three types of genes SARS-CoV-1, SARS-CoV-2 and H1N1 are taken. 39680 of genomic sequences of SARS-CoV-2 reported for Asian countries are collected from https://covidcg.org/, which is an open resource to track SNVs (single-nucleotide variations). For SARS-CoV-1, 54 mutated genomic sequences are considered. 35008 patients’ data of H1N1 type A are collected from NCBI influenza virus database (https://www.ncbi.nlm.nih.gov/genomes/FLU/Database/nph-select.cgi#mainform). The information about collected dataset are summarized at table.

### 3.1. Derive Degree of Mutation of each dataset

It is observed that mutations occurred at different positions of codon throughout the dataset. Here in this section we have tried to find out the highest occurrence of codon transitions. The degree of mutations for each mutation is analysed. It has been observed that mutations are majorly taken place of degree 3 for all datasets, which has been shown in figure1.

**Figure 1:**
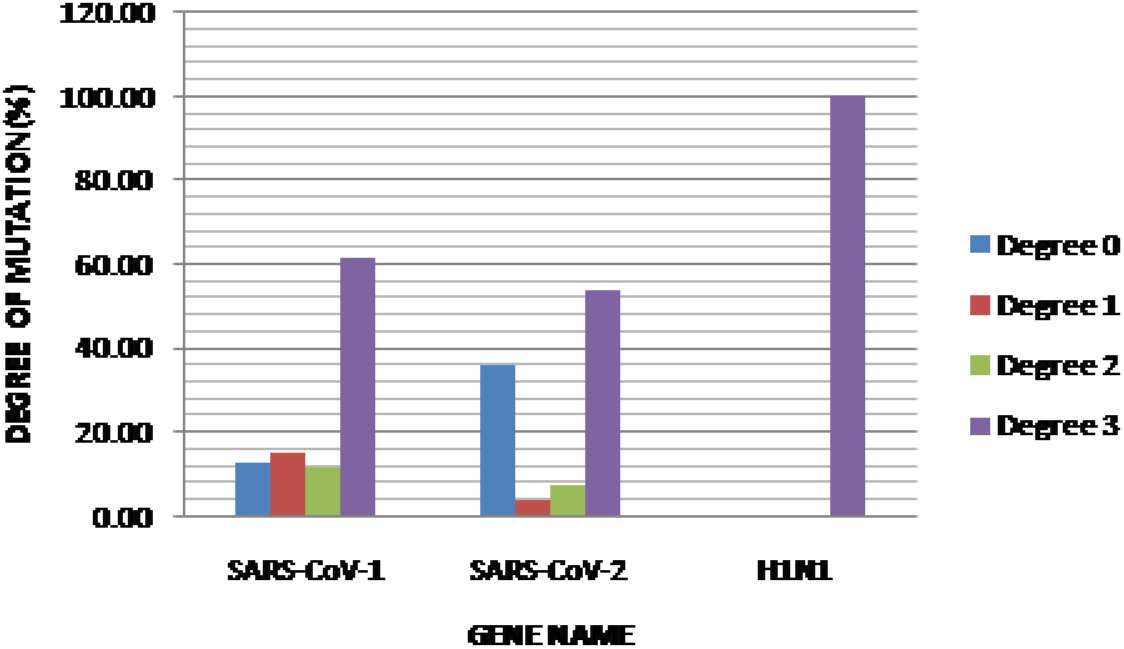
Percentage-wise analysis of degree of mutation

### 3.2. Position-wise analysis of mutations occur at 3 different nucleotide positions of codon

Codons are triplets comprising 3 nucleotides at its three positions. Mutations may occur at any of those three positions. In this sub section percentage wise calculations have been made on mutations occurred at those three different positions of codons for the SNVs of the 3 sets of genomic sequences taken. It has been observed that in the SNVs of SARS-CoV-1 (41.18%) the mutations majorly took place at 2nd positions. In the SNVs of H1N1 Type A mutations occurred in equal percentage at 1st and 2nd positions of codons. In SARS-CoV-2 (41.18%) the maximum mutations occurred at 1st positions of codon.

### 3.3. Representation of mutations occur in different codon positions based on the rules of cellular automata

Here in this subsection it is tried to make a mapping between rules defined by genetic code and 16 rules of Cellular Automata. The model is applied on the all three datasets taken. The Figure3 shows percentage of mutations occurred according to CA rule. It has been observed that the SNVs of SARS-CoV-2 has a trend to mostly follow the rules 4 (51%), whereas, CA rule 4 (19.01%) and CA rule 1 (18.71%) have approximately equal contributions in of SARS-CoV-1. The SNVs of H1N1 has the trend of rule 14 (51%). The rule 4 indicates the substitution of nucleotide C by T and rule 1 specifies substitution of T by C, i.e. between pyrimidines and rule 14 indicates the substitution of neucleotide G by A, i.e. between purines. Further microscopic view has been given on the codon position wise degree of mutations occurred in SARS-CoV-1 and SARS-CoV-2 where rule 4 (*C* → *T*) is applied and in H1N1 rule 14 (*G* → *A*) is applied maximum. The result is shown in Figure 4. It is remarkable that in both the datasets of SARS-CoV-1 and SARS-CoV-2 maximum mutations took place at 2nd position of codons and they are of degree 3. In H1N1 TYPE A virus all the mutations of degree 3 are taken places equally at the 1st and 2nd position of codons. Few transversions (15.29%) are also taken place in SARS-CoV-2. In these case base G of codons are substituted by T. In CA rule this substitution comes under rule 12.

**Figure 2:**
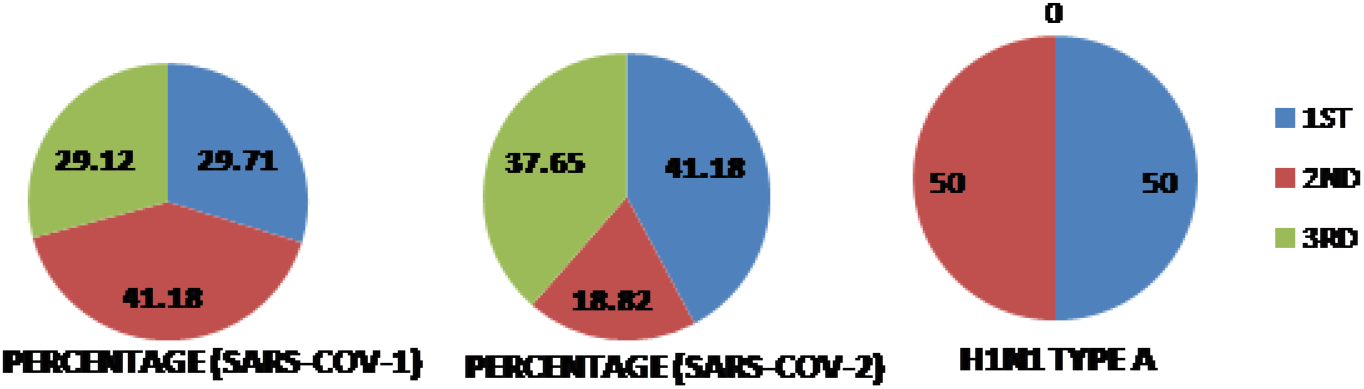
Representation of mutations occur in different positions of codon.

**Figure 3:**
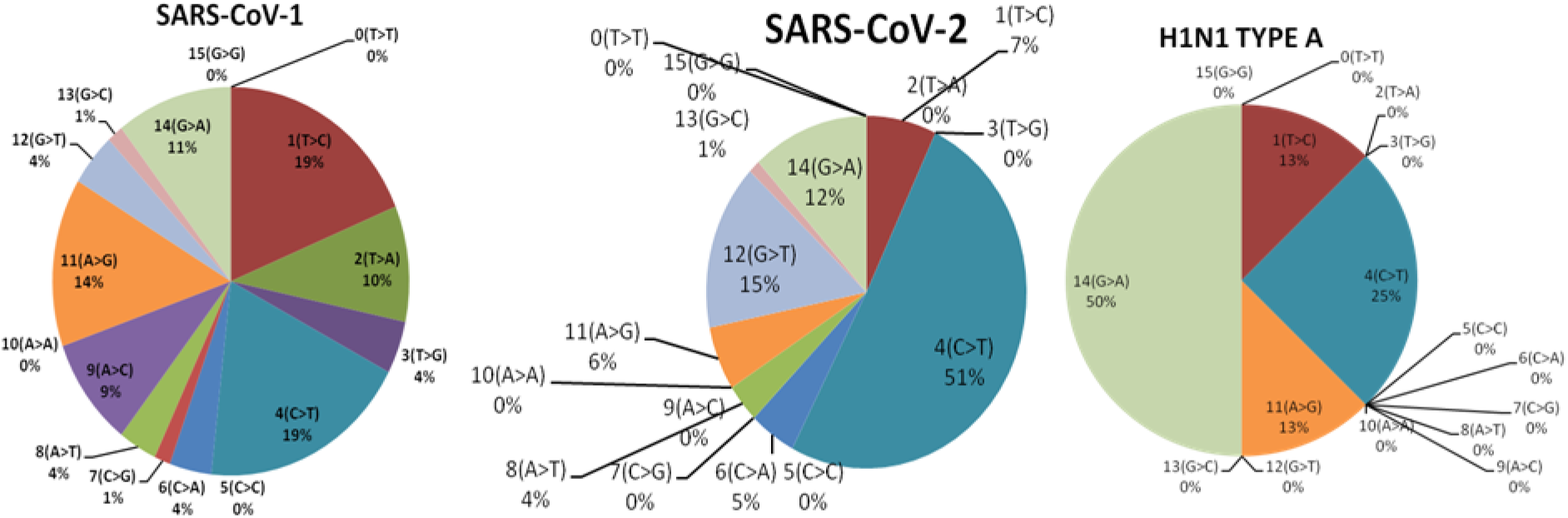
Percentage-wise mutations occur in three virus datasets according to 16 rules of cellular automata.

**Figure 4:**
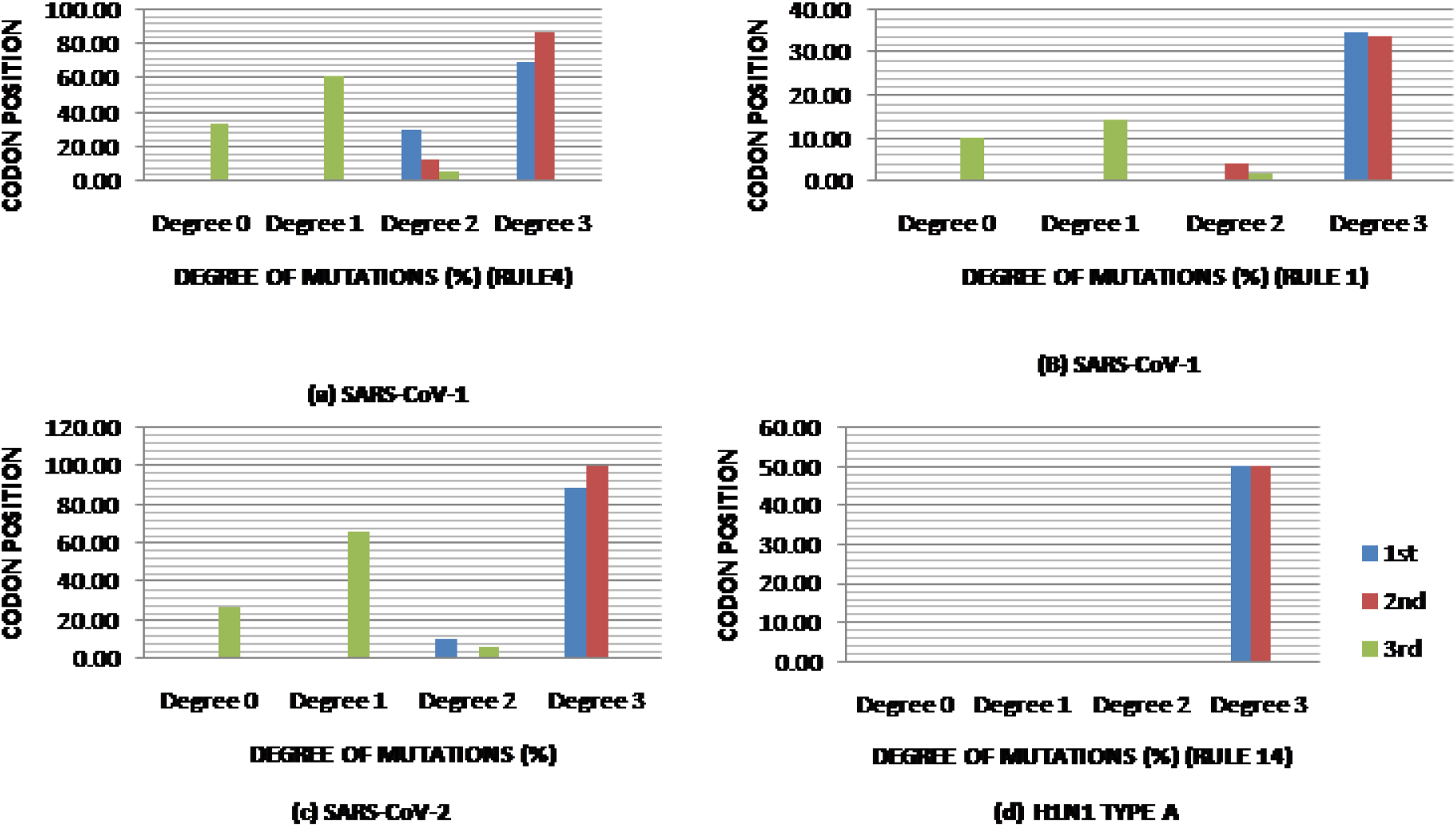
Codon position wise degree of mutations occurred in SARS-CoV-1 and SARS-CoV-2 where rule 4 is applied and in H1N1 where rule 14 is applied maximum. a) SARS-CoV-1 (b) SARS-CoV-2 (c) H1N1 type A.

Further analysis has been carried out with the mutations occurred under rule 4 for the datasets of SARS-CoV-1 and SARS-CoV-2 and under rule 14 for H1N1 Type A virus respectively (shown in Table 5. It has been found that some mutations have dominance over the others and codon position wise they have commonalities between SARS-CoV-1 and SARS-CoV-2. In both the datasets *L* → *L*, *L* → *F* are majorly found mutations at the first positions of codons and *T* → *I* at second positions. Individually frequently found mutations IN SARS-CoV-1 are *L* → *L*, *A* → *V*, *T* → *I*, *P* → *L*, *Y* → *Y*. In SARS-CoV-2 they are *L* → *L*, *F* → *F* and *Q* → *STOP CODON*. In H1N1 type A *S* → *N* and *H* → *Y* are found the most.

**Table 5:** Amino acid changes due to mutations occurred under rule 4 for the datasets of SARS-CoV-1 and SARS-CoV-2 and under rule 14 for H1N1 Type A virus

| Substitution | Codon Position | Possible AA Changes | SARS-CoV-1 | SARS-CoV-2 |
| --- | --- | --- | --- | --- |
| $C \rightarrow T$ | 1st | L→F, P→S, H→Y, R→W, R→C, Q→STOP CODON | H→Y, Q→STOP CODON, P→S, L→F, L→L | R→C, H→Y, L→F, L→L, P→S, Q→STOP CODON |
|  | 2nd | S→F, S→L, P→L, T→I, T→M, A→V | T→I, T→M, P→L, A→V, S→L, L→F | T→I, A→V, P→L, S→F |
|  | 3rd | Synonymous Changes (F→F, L→L, I→I, V→V, S→S, P→P, T→T, Y→Y, H→H, N→N, D→D, C→C, R→R, G→G) | N→N, T→T, S→S, I→I, Y→Y, L→L, D→D, A→A, G→G, V→V | Y→Y, C→C, F→F, H→H, I→I, L→L, N→N, S→S, T→T |
| Substitution | Codon Position | Possible AA Changes | H1N1 TYPE A |  |
| $G \rightarrow A$ | 1st | V→M, V→I, A→T, D→N, E→K, G→R, G→S | A→T, E→K | |
|  | 2nd | G→D, G→E, R→K, S→N, R→Q, R→H, W→STOP CODON, Y→C | S→N, G→E |  |
|  | 3rd | Synonymous Changes (L→L, S→S, STOP CODON→STOP CODON, L→L, P→P, Q→Q, R→R, T→T, K→K, V→V, A→A, E→E, G→G)<br>Non-synonymous Changes (M→I, W→STOP CODON) | None |  |

In SARS-CoV-1, few transversions are found, where substitutions are taken place between A and T, which are defined by CA rule 2 (*T* → *A*) and rule 8 (*A* → *T*). It is reported that the most harmful mutations due to substitutions take place between A and T. These kind of mutations change the hydropathy and polarity of amino acids. Hence, next point of investigation is carried out with it. It has been observed that according to CA rules, 9.61% and 3.51% of total SNVs found in the data set of SARS-CoV-1 are following the rule 8 (*T* → *A*) and rule 2 (*A* → *T*) respectively (shown in Table 6).

**Table 6:** Substitutions in SARS-CoV-1 according to CA rule 8 and rule 2.

| CA Rule (Substitution) | POSITION | AA CHANGED |
| --- | --- | --- |
| Rule 2 ($T \rightarrow A$) | 1st | STOP CODON→K, Y→N,<br>S→T, C→ S, L→I, L→M |
| | 2nd | I→K, V→E, M→K, I →N,<br>V→D, V→E, L→STOP<br>CODON, F→rY |
| | 3rd | I→I, Y→STOP CODON,<br>A→A, V→V, F→L |
| Rule 8 ($A \rightarrow T$) | 1st | N→Y, I→L, K→ STOP<br>CODON, I→F, N→Y,<br>STOP CODON→ L |
| | 2nd | STOP CODON→ L,<br>D→V, Q→L |
| | 3rd | G→G, A→A, L→F |

### 3.4. Discussion

In this article, point mutation (substitution type) of a codon has been associated with local transitions of Cellular Automata (CA) having 3-celled local configurations. Clearly, 16 substitutions are possible with four DNA bases A, C, T and G, which can be represented in terms of combinations of 16 local transition functions of CA. The experiment has been carried out with three sets of SNVs of three different viruses but the symptoms of the diseases caused by them are to some extent similar to each other. They are SARS-CoV-1, SARS-CoV-2 and H1N1 Type A viruses. The aim is to understand the impact of nucleotide substitutions in different codon positions on mutations occurred in a particular disease phenotype. With reference to the supplementary Table S1 it is to be noted that although the size of genomic sequences taken for all three viruses are huge, but H1N1 type A virus has comparatively very few variants. Codon usage bias is observed in all organisms even in viruses too. The reason behind may be either pressure of natural selection or due to biases in the mutation process. According to the origin and evolution theory of genetic code, codons are selected in such a way so that it can minimize the adverse effect of point mutations and translation errors. It has been observed that in all the datasets maximum mutations have taken place at the codons having degree of mutation 3. The codons having degree of mutation 3 are capable to change up to 3 amino acids due to substitution of nucleotides at a particular position. It has been observed that in the SNVs of SARS-CoV-1 the mutations majorly took place at 2nd positions but in SNVs of H1N1 type A 1st and 2nd positions of codons are equally affected. In SARS-CoV-2 the maximum mutations occurred at 1st positions of codon. The second position of codon is the most functionally constrained position and causes non-synonymous change. According to the nature of mutations, 16 CA rules of substitutions are classified into 3 classes namely, ‘No Mutation’, ‘Transition’ and ‘Transversion’. Experimental results find substitutions from CA class ‘Transition’ more than the other two classes. Transition mutations are more likely than transversions, because transversions make substitutions of nucleotides between purine (having 2 rings in it structure) and pyrimidine (having 1 ring). Hence, substitution of a single ring structure with another single ring structure is more likely than substitution of a double ring with a single ring. Transitions are more certain to change amino acids. Harmful substitutions from CA class ‘Transversion’ (rule 2 and rule 8) are noticed (13.16% in total) between bases A and T in some SNVs of SARS-CoV-1, which are responsible to make huge structural changes in existing proteins.

## 4. Conclusion

In this article a Cellular Automaton has been used to model substitutions of four DNA bases A, C, T and G at different positions of codons. Considering codon as a triplet, substitution of nucleotides may take place in any one of the three positions of a codon and cause point mutations. All possible point mutations have been represented here as functions of 16 CA transition rules. Point mutations may or may not make changes in the amino acids. The degree of mutation at a particular position of a codon defines the number of amino acids change due to substitution of nucleotides at each position of the codon, when other two positions of that codon are fixed. Hence, the degree of mutation specifies the capability of nucleotide substitutions in a particular position of a codon to produce new amino acids and their impacts in a particular disease pathogenesis. Thus, the aim of this work is investigating the codon alteration patterns due to nucleotide substitutions and their impact during mutations of a gene responsible for a particular disease. Hence, signature of a particular disease could be portrayed in the light of CA transition rules and codon alteration patterns.

## Supporting information

*Table S1.* Virus-wise specification of all the SNVs.

## References

[1] T. Négadi, The genetic code multiplet structure, in one number, arXiv preprint arXiv:0707.2011 (2007).

[2] Z. Zhou, Y. Dang, M. Zhou, L. Li, C.-h. Yu, J. Fu, S. Chen, Y. Liu, Codon usage is an important determinant of gene expression levels largely through its effects on transcription, Proceedings of the National Academy of Sciences 113 (2016) E6117–E6125.

[3] A. L. Labella, D. A. Opulente, J. L. Steenwyk, C. T. Hittinger, A. Rokas, Variation and selection on codon usage bias across an entire subphylum, PLoS genetics 15 (2019) e1008304.

[4] R. D. Knight, S. J. Freeland, L. F. Landweber, A simple model based on mutation and selection explains trends in codon and amino-acid usage and gc composition within and across genomes, Genome biology 2 (2001) 1–13.

[5] W. Huang, Y. Guo, N. Li, Y. Feng, L. Xiao, Codon usage analysis of zoonotic coronaviruses reveals lower adaptation to humans by sars-cov-2, Infection, Genetics and Evolution 89 (2021) 104736.

[6] S. Hormoz, Amino acid composition of proteins reduces deleterious impact of mutations, Scientific reports 3 (2013) 1–10.

[7] L. Bofkin, N. Goldman, Variation in evolutionary processes at different codon positions, Molecular Biology and Evolution 24 (2007) 513–521.

[8] P. Błażej, M. Wnętrzak, D. Mackiewicz, P. Mackiewicz, Correction: Optimization of the standard genetic code according to three codon positions using an evolutionary algorithm, Plos one 13 (2018) e0205450.

[9] A. Zeb, E. Alzahrani, V. S. Erturk, G. Zaman, Mathematical model for coronavirus disease 2019 (covid-19) containing isolation class, BioMed research international 2020 (2020).

[10] S. Jiang, Q. Li, C. Li, S. Liu, X. He, T. Wang, H. Li, C. Corpe, X. Zhang, J. Xu, et al., Mathematical models for devising the optimal sars-cov-2 strategy for eradication in china, south korea, and italy, Journal of translational medicine 18 (2020) 1–11.

[11] X. Wu, Y. Cai, X. Huang, X. Yu, L. Zhao, F. Wang, Q. Li, S. Gu, T. Xu, Y. Li, et al., Co-infection with sars-cov-2 and influenza a virus in patient with pneumonia, china, Emerging infectious diseases 26 (2020) 1324.

[12] D. Silvestro, A. Antonelli, N. Salamin, T. B. Quental, The role of clade competition in the diversification of north american canids, Proceedings of the National Academy of Sciences 112 (2015) 8684–8689.

[13] A. Sengupta, S. S. Hassan, P. P. Choudhury, Clade gr and clade gh isolates of sars-cov-2 in asia show highest amount of snps, Infection, Genetics and Evolution 89 (2021) 104724.

[14] J. Kari, Theory of cellular automata: A survey, Theoretical computer science 334 (2005) 3–33.

[15] P. Kiran Sree, I. R. Babu, S. Usha Devi N, Cellular automata and its applications in bioinformatics: A review, arXiv e-prints (2014) arXiv-1404.

[16] C. Burks, D. Farmer, Towards modeling dna sequences as automata, Physica D: nonlinear phenomena 10 (1984) 157–167.

[17] S. GCh, I. Karafyllidis, M. Ch, V. Mardiris, A. Thanailakis, P. Tsalides, A cellular automaton model for the study of dna sequence evolution., Computers in Biology and Medicine 33 (2003) 439–453.

[18] M. Ch, S. GCh, V. Mardiris, I. Karafyllidis, N. Glykos, R. Sandaltzopoulos, Reconstruction of dna sequences using genetic algorithms and cellular automata: Towards mutation prediction?, Biosystems 92 (2008) 61–68.

[19] P. P. Chaudhuri, S. Ghosh, A. Dutta, S. P. Choudhury, A New Kind of Computational Biology: Cellular Automata Based Models for Genomics and Proteomics, Springer, 2018.

[20] R. Dogaru, L. O. Chua, Mutations of the” game of life”: A generalized cellular automata perspective of complex adaptive systems, International Journal of Bifurcation and Chaos 10 (2000) 1821–1866.

[21] A. Madain, A. L. A. Dalhoum, A. Sleit, Application of local rules and cellular automata in representing protein translation and enhancing protein folding approximation, Progress in Artificial Intelligence 7 (2018) 225–235.

[22] S. Ghosh, S. Bhattacharya, Computational model on covid-19 pandemic using probabilistic cellular automata, arXiv preprint arXiv:2006.11270 (2020).

[23] S. Basu, S. Ghosh, Fuzzy cellular automata model for discrete dynamical system representing spread of mers and covid-19 virus, in: Internet of Medical Things for Smart Healthcare, Springer, 2020, pp. 267–304.

[24] P. K. Sree, S. U. D. Nedunuri, A novel cellular automata classifier for covid-19 trend prediction, Journal of Health Sciences 10 (2020).

[25] C. Beauchemin, J. Samuel, J. Tuszynski, A simple cellular automaton model for influenza a viral infections, Journal of theoretical biology 232 (2005) 223–234.

[26] C. Beauchemin, Probing the effects of the well-mixed assumption on viral infection dynamics, Journal of theoretical biology 242 (2006) 464–477.

[27] C. Beauchemin, S. Forrest, F. T. Koster, Modeling influenza viral dynamics in tissue, in: International Conference on Artificial Immune Systems, Springer, 2006, pp. 23–36.

[28] F. H. Crick, The origin of the genetic code, Journal of molecular biology 38 (1968) 367–379.

[29] S. Ghosh, S. Basu, Some algebraic properties of linear synchronous cellular automata, arXiv preprint arXiv:1708.09751 (2017).

[30] S. Ghosh, Evolutions of some one-dimensional homogeneous cellular automata, Complex Systems 30 (2021).

